# One year of gluten free diet impacts gut function and microbiome in celiac disease

**DOI:** 10.1101/2024.06.20.599876

**Authors:** Carolyn M. Costigan, Frederick J. Warren, Anthony P. Duncan, Caroline L. Hoad, Nina Lewis, Trevor Hill, Colin J. Crooks, Paul S. Morgan, Carolina Ciacci, Paola Iovino, David S. Sanders, Falk Hildebrand, Penny A. Gowland, Robin C. Spiller, Luca Marciani

**Affiliations:** Nottingham Digestive Diseases Centre, Translational Medical Sciences, School of Medicine, University of Nottingham, Nottingham NG7 2UH, United Kingdom; NIHR Nottingham Biomedical Research Centre, Nottingham University Hospitals NHS Trust, Nottingham NG7 2UH, United Kingdom; Quadram Institute Bioscience, Norwich Research Park, Norwich, NR4 7UQ, United Kingdom; Digital Biology, Earlham Institute, Norwich Research Park, Norwich, Norfolk, NR4 7UQ, United Kingdom; Sir Peter Mansfield Imaging Centre, School of Physics and Astronomy, University of Nottingham, Nottingham NG7 2RD, United Kingdom; Nottingham University Hospitals NHS Trust, Nottingham NG7 2UH, United Kingdom; Medical Physics, School of Medicine, University of Nottingham NG7 2UH, Nottingham, United Kingdom; Department of Medicine and Surgery, University of Salerno, 84081 Salerno, Italy; Department of Clinical Medicine, School of Medicine and Population Health, University of Sheffield, Sheffield S10 2HQ, United Kingdom

**Author notes:** CMC and FJW are joint First Authors.

**Keywords:** Celiac disease, microbiome, gluten free diet, MRI

## Abstract

**BACKGROUND & AIMS:** Currently, the main treatment option for coeliac disease (CD) is a gluten free diet (GFD). This observational cohort study investigated the impact of CD and 1 year of GFD on gut function and microbiome.

**METHODS:** 36 newly diagnosed patients and 36 healthy volunteers (HV) were studied at baseline and at 12 months follow up. Small bowel water content (SBWC), whole gut transit time (WGTT) and colon volumes were measured by MRI. Stool samples DNA was subjected to shotgun metagenomic sequencing. Species level abundances and gene functions, including carbohydrate active enzymes (CAZymes) were determined.

**RESULTS:** SBWC was significantly higher in people with CD 157±15 mL versus HVs 100±12 mL (p=0.003). WGTT was delayed in people with CD 68±8 hours versus HVs 41±5 hours (p=0.002). The differences reduced after 12 months of GFD but not significantly. Wellbeing in the CD group significantly improved after GFD but did not recover to control values. CD faecal microbiota showed high abundance of proteolytic gene functions, associated with *Escherichia coli, Enterobacter* and *Peptostreptococcus*. GFD significantly reduced *Bifidobacteria* and increased *Blautia wexerelae*. Microbiome composition correlated positively with WGTT, colonic volume and *Akkermansia municphilia* but negatively with *B.wexerelae*. Following GFD the reduction in WGTT and colonic volume significantly associated with increased abundance of *B.wexerelae*. There were also significant alterations in CAZyme profiles, specifically starch and arabinoxylan degrading families.

**CONCLUSIONS:** CD impacted gut function and microbiota. GFD ameliorated but did not reverse these effects, significantly reducing *Bifidobacteria* associated with reduced intake of resistant starch and arabinoxylan from wheat.

---

Coeliac disease (CD) requires a life-long commitment to a gluten free diet (GFD) treatment^1,2^. The aim of GFD is recovery of the small bowel mucosa and reversal of the enteropathy. Despite a long-term adherence to GFD, a significant proportion of people with CD report persistent gastrointestinal (GI) symptoms^3^.

Small bowel water content (SBWC) in untreated patients with CD measured using magnetic resonance imaging (MRI) was increased, a feature hypothesised to be due to a combination of impaired motility and absorption.^4^ Manometric, breath test and camera pill studies described an underlying gastrointestinal dysmotility with prolonged oro-cecal transit times.^5^ Such GI motor disorders tended to resolve on GFD^6^ suggesting that the motor dysfunction may be related to mucosal inflammation.^7^

CD also impacts the gut microbiome, with reported increased abundance of Firmicutes and Proteobacteria, and reductions in beneficial *Bifidobacterium*.^8^ Studies in CD have identified higher levels of *Escherischia coli,* Clostridiaceae and Enterobacteriaceae, and lower *Bifidobacterium* in the duodenal mucosa.^9, 10^ The GFD tends to have a lower dietary fibre intake than habitual diets^11^ and may have effects on the gut microbiome independent of CD. In healthy volunteers GFD reduced the abundance of *Bifidobacterium* (specifically *Bifidobacterium longum*) and *Lactobacillus*, and increased the abundance of Enterobacteriaceae (specifically *E.coli*).^12^ There may also be a difference in the metabolic activity of intestinal microbial flora in children with CD compared to those without the disease, and this difference may not be affected by diet treatment.^13^

The reasons for alterations in the gut microbiome of CD patients and correlations with GI function and symptoms are not clearly understood. This study aimed to investigate the small bowel water content, gut transit time, colon volumes, symptoms and microbiome of CD patients at diagnosis, and the impact of following one year of GFD using combined non-invasive MRI measurements with microbiome analysis using shotgun metagenomic approaches. The primary hypothesis was that GFD will reduce the fasting SBWC.

## Methods

### Study population

Thirty-six patients newly diagnosed with CD were prospectively recruited before starting treatment with a GFD. The patients commenced their GFD treatment immediately after their baseline study visit. Adherence was measured by repeating the coeliac serology testing at 12 month follow up, as well as using Biagi scores.^14^ The 12 months follow up group included patients providing a stool sample and having GFD diet adherence confirmed by TTG blood sample and Biagi score ≥3.

A parallel group of 36 HVs frequency matched for age and sex were prospectively recruited for comparison data. They were not following a GFD diet. Both cohorts were scanned twice and approximately 12 months apart, to investigate temporal effects in the endpoints measured and assess the impact of any seasonal change. The 12 months follow up group included healthy participants who did not exclude gluten from the diet, provided a stool sample and had confirmed negative CD screening blood sample.

Patient eligibility criteria also included: being 18-70 years old; on a gluten-containing diet; IgA-TG2 or IgA-DGP, or IgG-DGP, and duodenal biopsy confirming CD; recruited within a month of their duodenal biopsy. The healthy volunteers were required to have a screening blood sample negative for CD and to have no co-morbidities. All participants were not to have had any antibiotic or probiotic treatment in the 4 weeks preceding the study days.

### Study protocol

Participants recorded on Bristol Stool Form Scales time and consistency of every bowel movement for the seven days prior to each study day. On each study day the participants were asked to provide a stool sample using a supplied collection kit which included aliquot laboratory tubes and a biosafety container with freezer packs, to be kept in the home freezer (-5°C) until needed for sample collection. The container was then brought to the study unit and the samples were then stored immediately at -80°C until they underwent DNA extraction, sequencing to a depth of ∼10GB per sample and metagenomic processing.

At 8am the day before the study day, the participants were instructed to swallow, before their usual breakfast, 5 inert plastic MRI transit capsule markers to measure WGTT.^15^ They were also asked to fast from their evening meal, eaten by 8pm, until the following morning study, with only water allowed until bedtime. In the morning, fasted, MRI scans were acquired to measure SBWC, colonic volumes and WGTT.

Fasting breath hydrogen (H_2_) was measured using a portable breath H_2_ meter (Gastro+Gastrolyzer, Bedfont Scientific, Kent, UK). Psychometric assessment was performed using the Hospital Anxiety and Depression Scale (HADs) and the Personal Health Questionnaire (PHQ-15), also scored separately for the three GI symptoms (PHQ-3). GI symptom intensity on each study day was also measured using visual analogue scales (0–100 mm VAS) for abdominal bloating, flatulence, nausea and abdominal pain.

The study was registered on ClinicalTrials.gov (NCT02551289).

### Statistical and bioinformatics analysis

There was no data available to estimate the size of the change in SBWC in patients after a GFD diet. Inference was drawn from previous MRI data ^4^ in n=20 untreated, newly diagnosed patients with CD showed fasting WBWC of 202 ml±(SD)115 ml. From this it was predicted that we could detect a change of 40% (a reduction of 80 ml volume) after gluten free diet with a power of 90% and a Type I error probability of 0.05 using n=24 patients in a paired study design. This would be considered a clinically significant change after GFD; this change reflects twice the level of normal control values of 65 ± 43 ml volume.^4^ A recruitment target of n=36 was planned to allow for dropouts and to increase power for secondary outcomes.

All analyses, both descriptive and statistical were carried out in Stata MP4 v18.0 (StataCorp LLC). Normality of data was assessed via histogram with the normal distribution overlaid. For comparing outcomes between the study groups, differences between mean values at baseline, and mean changes from baseline to follow-up were compared via unpaired t-test. Where the normality of the differences was in question, the analysis was repeated using the unpaired two-samples Wilcoxon test. Statistical tests were two-sided. A p-value of 0.05 was used to assess statistical significance. All data are presented as mean±SEM.

LEfSe (Linear discriminant analysis Effect Size)^16^ was used to identify individual taxa and functionally annotated genes and metabolic pathways. A p-value of 0.05 was used as a cut-off for statistical significance, and an LDA value of 2.5 was used to identify discriminating features. Linear mixed effects modelling using the MaAsLin2 package^17^ was used to identify correlations between individual taxa and functionally annotated genes and metabolic pathways.

All authors had access to the study data and reviewed and approved the final manuscript.

## RESULTS

### Baseline characteristics and differences between groups

The participants were 36 patients newly diagnosed with CD and 36 healthy controls. They were predominantly middle-aged females with average BMI slightly above the healthy range. At baseline 21 patients had a GFD compliance Biagi score of 0 and 15 of them had a score of 1 indicating that at baseline none of them was following a GFD.^18^

The study procedures and visits were well tolerated by the participants in both groups with few dropouts (see CONSORT diagrams in Supplementary Figure 1). The two cohorts were matched for age, sex and body mass index as indicated in **Table 1**.

**Table 1.**
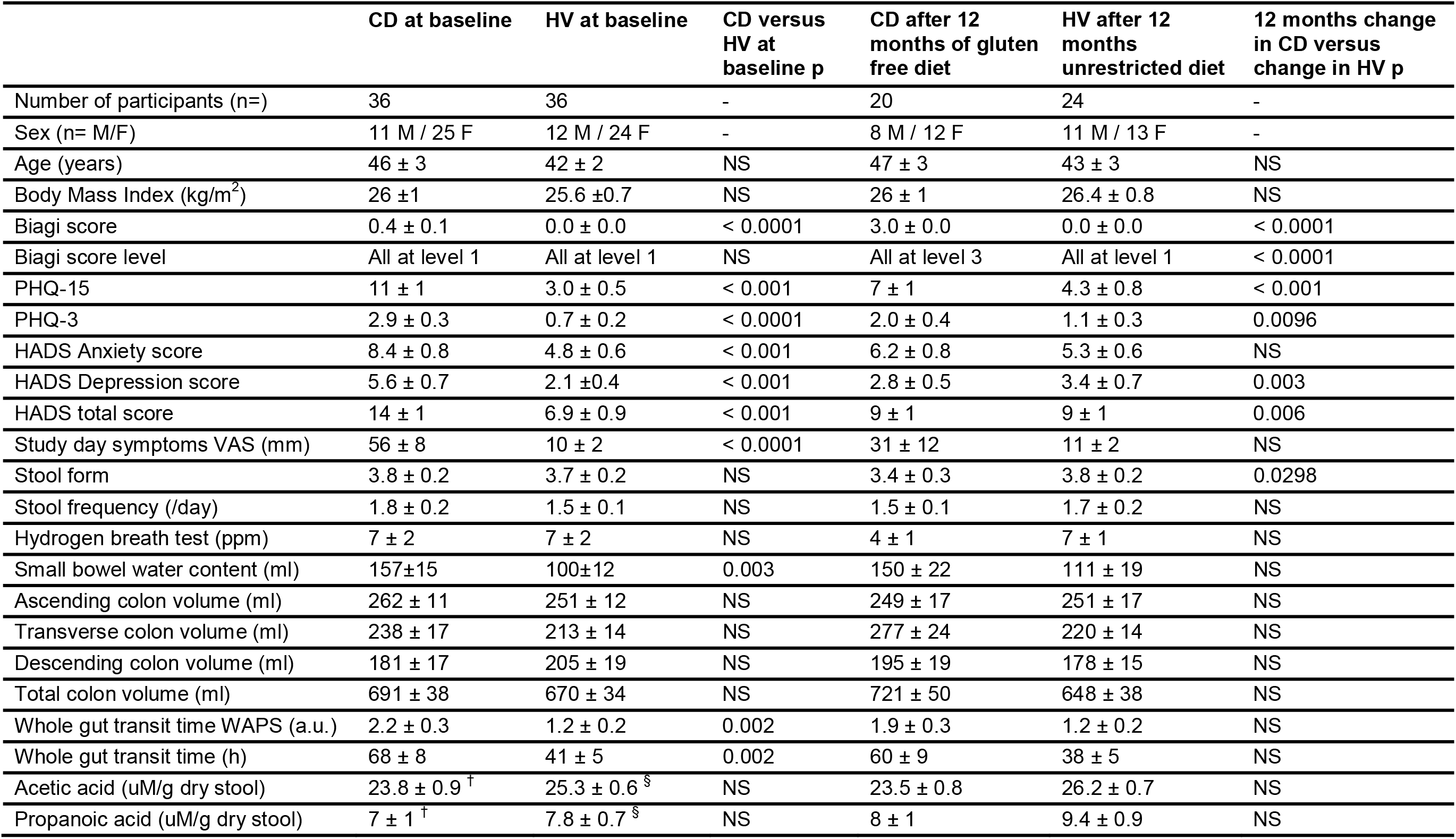
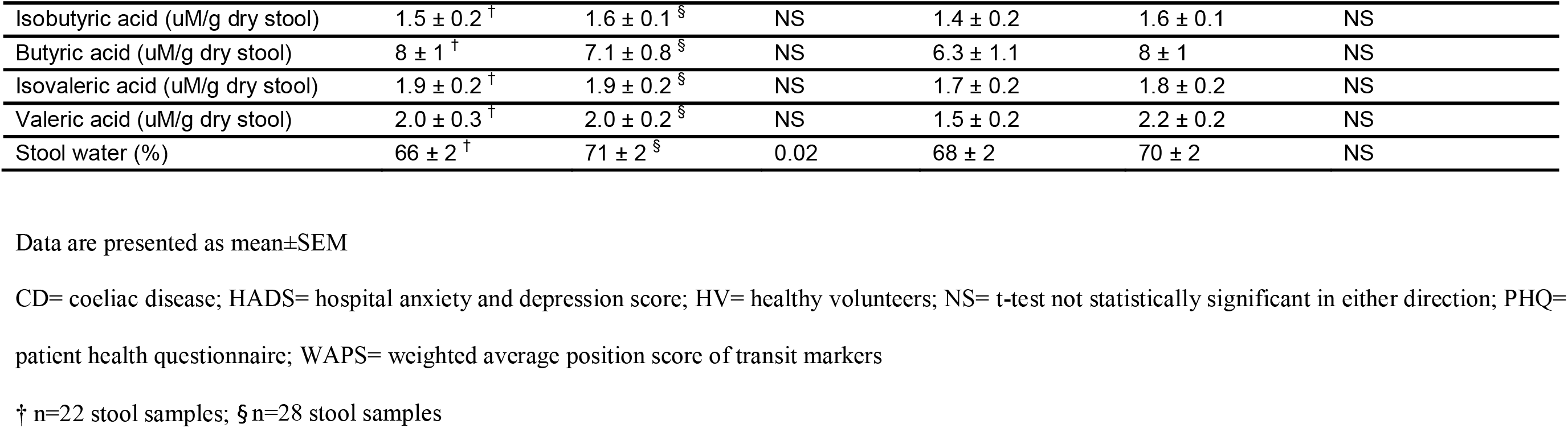
Demographic, behavioral and gastrointestinal endpoints at baseline and at 12 months follow up.

Patients with CD had higher levels of somatisation at baseline assessment, compared to the HVs as shown by the patient health questionnaire (PHQ-15) and in the subset of questions relating to GI problems PHQ-3 (see also Supplementary Figure 2). They also had higher scores for depression and anxiety and reported higher levels of GI symptoms on the visual analogue scale (VAS) on the MRI study day. People with CD had no significant differences in bowel habits and SCFAs to the healthy volunteers but had 5% lower percentage water in the stools.

At the baseline MRI assessment, the SBWC of the people with CD was 57% higher than that of the HCs (Figure 1A, p=0.003). Baseline whole gut transit time (WGTT) measured using MRI transit markers was 83% longer in people with CD compared to healthy volunteers (Figure 1B, p=0.002).

**Figure 1.**
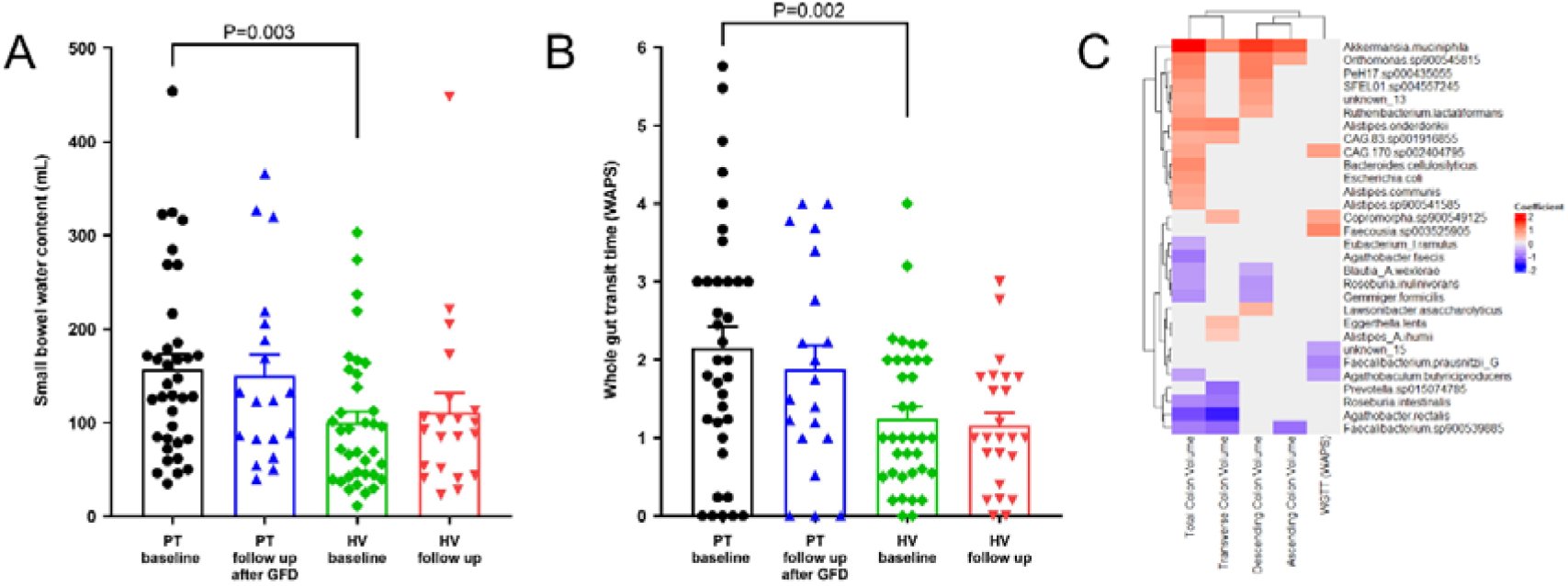
(A) Small bowel water content (B) Whole gut transit time (C) Multivariate regression analysis showing bacterial species which are significantly associated with MRI markers of WGTT and colonic volumes. Red colour indicates positive association, and blue negative association. Data are presented as mean±SEM.

Gut transit and colonic volumes may impact gut microbiome composition.^19^ Here whole gut transit time and colonic volumes were found to correlate with several microbial species (Figure 1C). Slower gut transit was correlated with *Akkermansia municiphila*, several species of the genus *Alistipes* and *Bacteroides* and the species *Ruthenibacterium lactatiformans,* while faster gut transit correlated with *Faecalibacterium prausnitzii, Gemmiger formicilis, Blautia wexlerae* and several *Agathobacter* species. These findings are consistent with recent studies which have associated these microbial groups with gut transit time using alternative methods to determine WGTT (e.g. stool consistency or coloured dyes)^19, 20^. Species level associations with other patient measures are shown in Supplementary Figures 3-5.

At baseline, prior to starting a GFD, while there were not major community level differences in microbiome composition (p>0.05, PERMANOVA) (Figure 2A), there were significant differences in individual taxa observed between the HV and CD cohorts (Figure 2B). The microbiomes in CD were significantly enriched in bacterial taxa including Oscillospiraceae and Peptostreptococcales compared to the HVs. Conversely, the HVs had higher abundances of the genus *Rumminococcus_D* and the species *B. wexlerae*. It is interesting to note that *B.*wexlerae was associated with fast gut transit times and smaller colonic volumes (Figure 1), so the higher abundance of *B.wexlerae* in healthy volunteers at baseline may reflect differences in passage time and colon volume between HV and people with CD. At baseline relatively few differences in CAZyme composition (Figure 2C) were observed between the healthy cohort and CD, although higher levels of GH20 and GH33 were observed in CD, both enzyme families involved in host glycan metabolism, and CBM5, involved in chitin metabolism. Several metabolic pathways were found to be elevated in CD, many of which are related to protein metabolism. This may reflect malabsorption of protein in the upper-GI tract of patients with CD.

**Figure 2.**
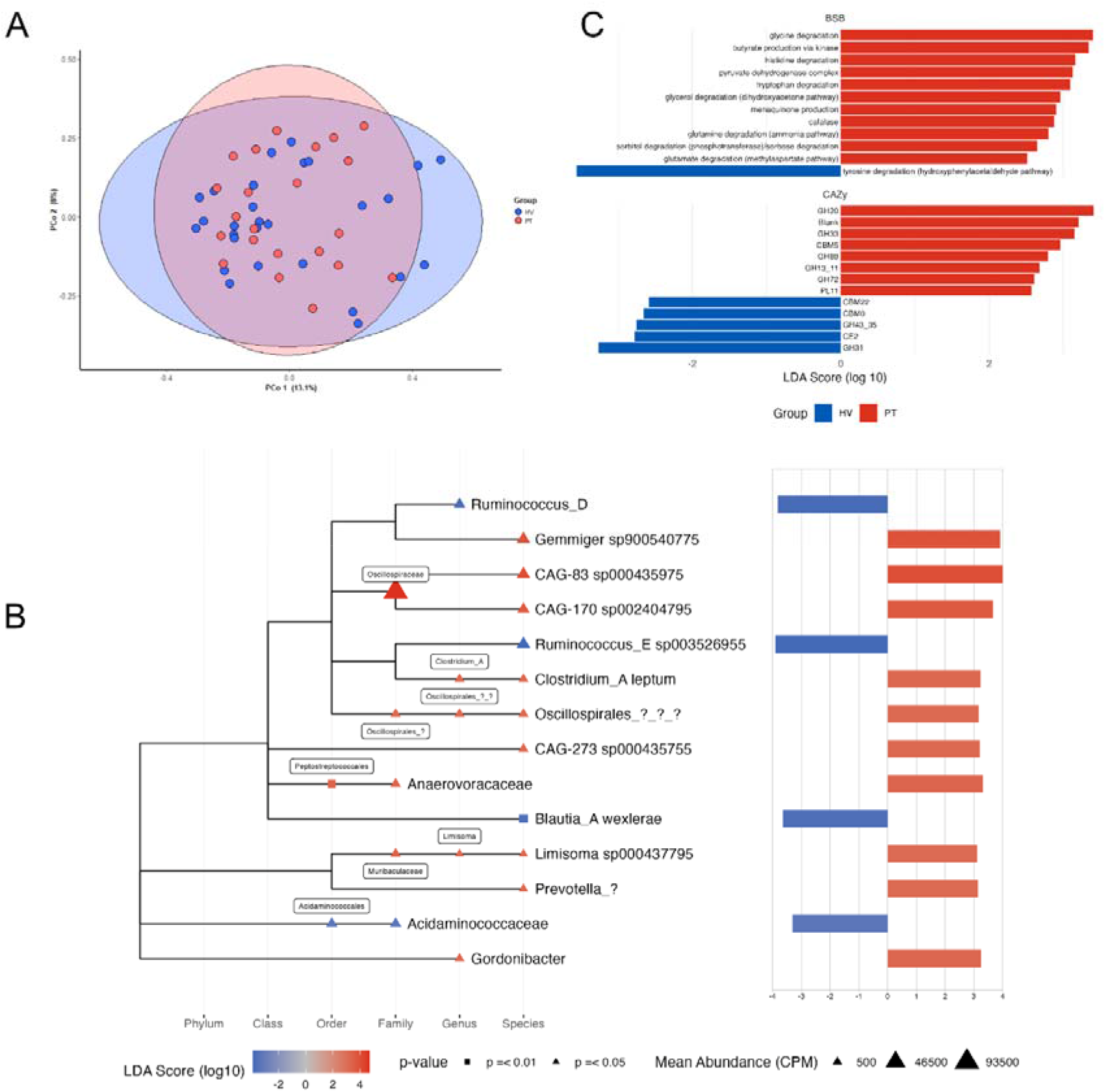
Baseline microbiome composition. (A) PCoA plot of Bray-Curtis distances of species diversity coloured by HV vs. PT (PERMANOVA p>0.05). (B) Differentially abundant microbial taxa identified by the LEFse algorithm (p=0.05) between HV and PT at baseline. (C) Differentially abundant CAZymes and BSB gene functional pathways (p=0.05). A blue bar indicates enrichment in HV and a red bar indicates enrichment in the PT group.

### Effects of gluten free diet

At 12 months follow up all the patients included in the data had achieved a Biagi score of 3 indicating that they followed a strict GFD (Biagi level 3). After 12 months of GFD the symptoms and wellbeing of the group of patients with CD had significantly improved compared to baseline values against the HVs’ seasonal variation although most of the endpoints had not recovered to HV values. The somatization scores in people with CD had significantly reduced by 36% for the PHQ-15 questionnaire and by 31% in the PHQ-3 subset of GI problems. Their total scores for depression and anxiety reduced by 36% and the VAS GI symptoms reduced by 45%. Stool form in people with CD increased on average by half a point on the Bristol stool scale and the percentage water in their stools increased by 2%. The higher amount of SBWC detected at baseline did not reduce after GFD. Whole gut transit time reduced by 14% after GFD but the change was modest and it was not significant compared to HVs.

Following 12 months of GFD did not result in community level shifts in the gut microbiome composition of patients with CD (Figure 3A) but did result in changes in individual taxa and metabolic taxa which were distinct from the baseline differences between patients with CD and HVs. Whilst no significant differences were observed between baseline and 12-month follow up for the healthy volunteers HVs, several differences were observed in the patients with CD. There was an increase in abundance of *Blautia_A*, which may be related to the improvements in WGTT. A significant increase in mucin degradation pathways was also observed which may reflect changes in WGTT. There were also increases in abundance in *Bacteroides uniformis*. Notably, there was a significant reduction in Actinomycetales, the class of bacteria encompassing Bifidobacterial species in patients with CD on the GFD. This was associated with a reduction in the *Bifidobacterium* specific metabolic pathway (Bifid shunt), suggesting that a GFD may have specific impacts on Bifidobacterial abundances.

**Figure 3.**
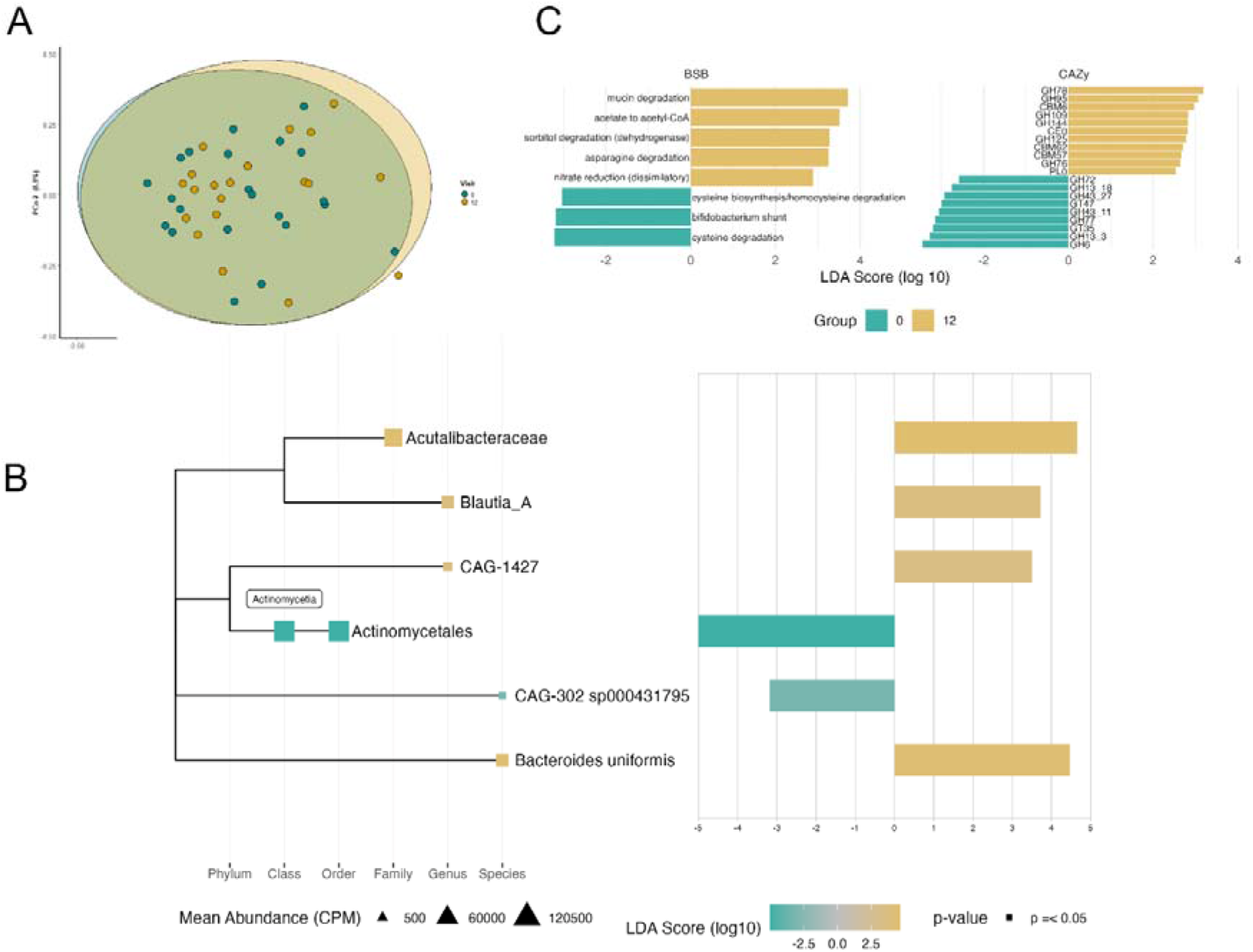
Gut microbiome changes from baseline to follow-up. (A) PCoA plot of Bray-Curtis distances of species diversity coloured by baseline vs. follow-up (PERMANOVA p>0.05). (B) Differentially abundant microbial taxa identified by the LEFse algorithm (p=0.05) between baseline and follow-up in CD patients. (C) Differentially abundant CAZymes and BSB gene functional pathways (p=0.05). A teal bar indicates enrichment at baseline and a mustard bar indicates enrichment at follow-up.

At the 12-month follow-up there was a significant difference between the HVs and the patients with CD in microbiome composition (p=0.002, F-statistic=2.1943, PERMANOVA) (Figure 4A). The reduction in *Bifidobacterium* observed as a result of GFD can clearly be observed in the CD group compared to the HVs at follow-up (Figure 4B). Specifically, the abundance of *Bifidobacterium longum* and *Bifidobacterium breve* is lower in CD patients than HVs. Additionally, *E.coli*, *Bacteriodes ovatus, Alistipes communis, Roseburia hominis* and several other taxa were elevated in CD compared to HVs.

**Figure 4.**
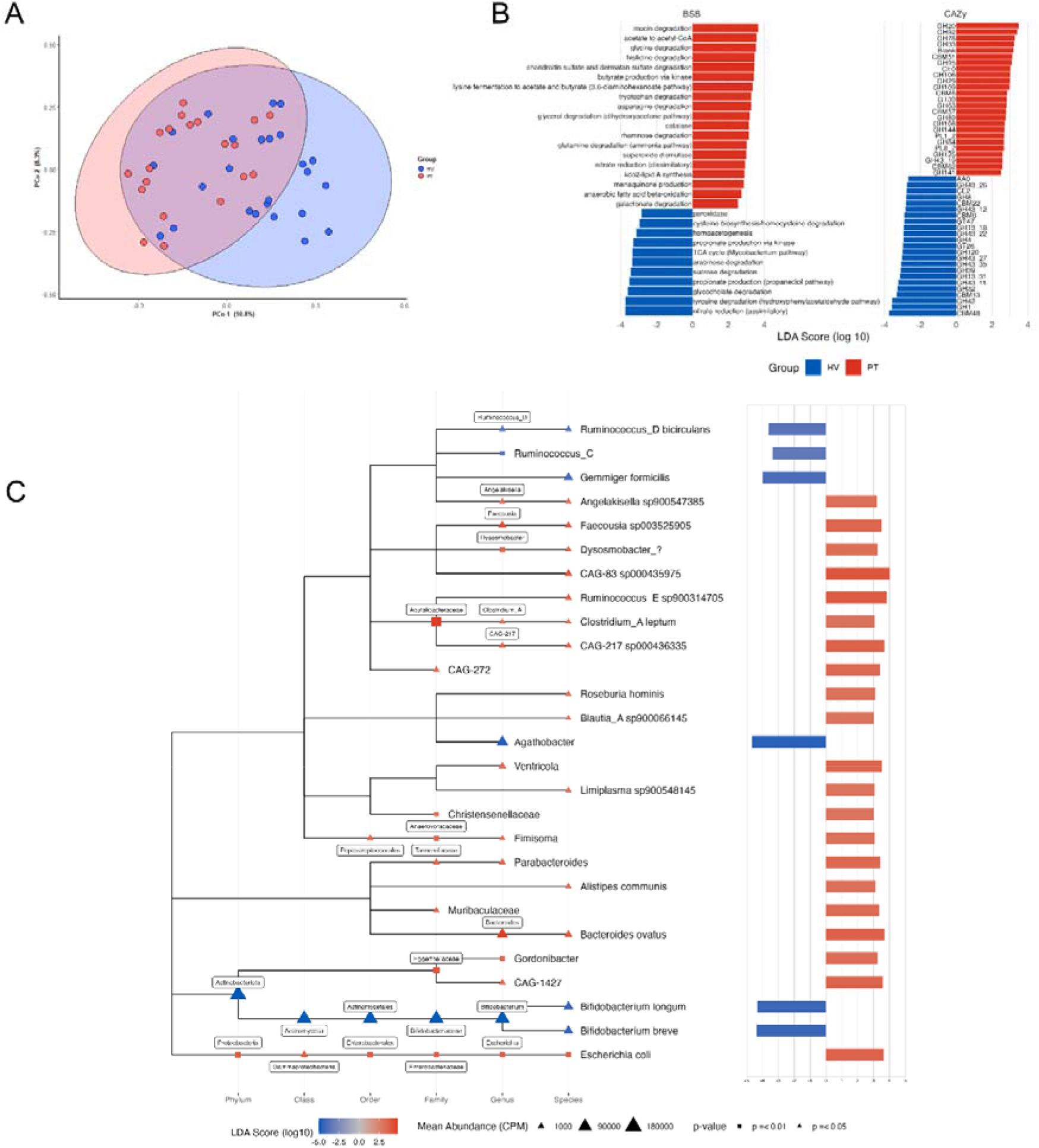
12 months follow-up microbiome composition. (A) PCoA plot of Bray-Curtis distances of species diversity coloured by HV vs. PT (PERMANOVA p=0.002, F-statistic=2.1943). (B) Differentially abundant microbial taxa identified by the LEFse algorithm (p=0.05) between HV and PT at baseline. (C) Results of LEFse analysis identifying differentially abundant CAZymes and BSB gene functional pathways (p=0.05). A blue bar indicates enrichment in HV and a red bar indicates enrichment in the PT group.

As well as differences in microbial community composition between the people with CD and HVs alterations in microbial metabolic pathway abundances were observed (Figures 4B and 4C) in addition to the differences observed at baseline. These may be related to the removal of the two major dietary fibre constituents in wheat, i.e. resistant starch and arabinoxylan caused by the GFD. The HVs were found to have significantly higher arabinose and sucrose degradation pathways, related to arabinoxylan and starch metabolism, respectively. The differences in CAZyme profile between HVs and patients also provides evidence for this. The CAZyme profile of healthy volunteers is enriched in several GH13 sub-families and CBM48 which are involved in starch metabolism, and in GT47 and several sub-families of GH43, which comprise xylanase and arabinofuranosidase enzymes involved in arabinoxylan metabolism. These changes in microbiome composition are also associated with shifts in the enterosignature^21^ groups between the HV and PT groups (Supplementary Figures 6 and 7). While at baseline there were no significant differences observed in enterosignatures between the PT and HV groups, at follow-up there was significantly higher relative abundance of the ES-Esch (comprising contributions from the genera *Escherichia, Citrobacter, Enterobacter, Klebsiella and Staphylococcus^21^*). There was also a significant reduction in the ES_Bifi enterosignature group relative abundance (Supplementary Figure 7B), specifically in the PT group compared to the HV group, reflecting the changes observed in abundance of *B. longum* and *B.breve* in Figure 4 at a community level.

Of note, *B. longum* is recognised as a keystone degrader of arabinoxylan^22^ and harbours a significant number of genes encoding GH43 enzymes in its genome and has been recognised to increase in abundance in response to dietary wheat arabinoxylan supplementation.^23^ In Supplementary Figure 8 the CAZYme profiles of *Bifidobacterium* MAGs from this study are shown. The genome of *B.longum* contained 14 copies of GH43 genes, more than any other Bifidobacterial genome identified in this study. In addition, the *B.breve* genome contains high numbers of gene copies of GH13 and CBM48, related to resistant starch degradation.

## Discussion

This study provided a detailed picture of the shifts in microbiome composition and function that occur in patients with CD at diagnosis and following 1 year of GFD compared to matched HVs. In common with previous studies we did not identify a ‘CD microbiome’ signature.^24^ We did, however, identify significant differences in individual taxa and metabolic pathways which may be associated with metabolism of excess malabsorbed protein reaching the gut and inflammation, as has been hypothesised to occur in CD.^24^

Following GFD for 1 year resulted in further alterations to the gut microbiome and significant community level differences between healthy volunteers and patients. Several taxa including *B.longum*, *B.breve* and *R.bicirculans* were lower in patients with CD compared to HVs, and conversely *E.coli* abundance was higher.^8, 25^ Studies which have investigated the microbiome changes in HVs following a GFD have identified similar changes.^12, 26^ In this study, we further highlight changes in enterosignatures in patents with CD following GFD.^21^

The functional microbiome analysis indicated alterations in carbohydrate metabolism, specifically relating to the two major sources of dietary fibre in wheat, arabinoxylan and resistant starch.^27^ *B.longum* strains have been characterised as being keystone arabinoxylan degraders^22^ and are increased in abundance in response to dietary interventions with arabinoxylan.^23^ Therefore, the changes observed in gut microbiome of patients with CD may result from a complex interaction of factors, including dietary as a result of following GFD and changes in gut physiology as a consequence of CD.

The SBWC was increased in the patients with CD at baseline compared to the HVs, but this showed no correlation with microbiome composition. Increased SBWC in CD may reflect the net effect of impaired absorption associated with villous atrophy and increased secretion associated with crypt hyperplasia, and also impaired motility. In this study no differences in colon volumes. ^4^ WGTT was significantly delayed in the CD cohort at baseline and although this improved after GFD it was still slower than in the healthy volunteers at follow up. Delayed WGTT in untreated CD may be due to a number of inter-related factors such as mucosal damage and inflammation affecting gut motility, malabsorption of food constituents and gut hormone derangement.

In this study, different microbial species correlated with markers of gut environment^21, 22^. Some of the gut microbiome shifts associated with GFD may therefore be related to alterations in transit.^28^

GI symptoms and quality of life reported by the newly diagnosed patients were poorer than that in the healthy volunteers. Although some of these measures improved after GFD treatment, their wellbeing and symptoms were still worse overall than the HVs at follow up.

In conclusion, the present study shows that gut function and microbiome are impacted by CD. One year of GFD treatment did not reverse the abnormalities and had a negative impact on the microbiome, with reduction of bifidobacterial abundance, increase in proteolytic species and reductions in starch-degrading and arabinoxylan-degrading CAZyme families. This potentially opens the possibility of developing interventions to reverse the negative impact of GFD with targeted prebiotic and/or symbiotic products.^29^

## Correspondence

Address correspondence to: Professor Luca Marciani, PhD, Nottingham Digestive Diseases Centre, Translational Medical Sciences, School of Medicine, University of Nottingham, Nottingham, NG7 2UH, United Kingdom.

[BACK]

## Acknowledgements

The authors sincerely thank all participants who have contributed to this study. The authors would also like to thank J. Price (Nottingham University Hospitals NHS Trust), specialist coeliac dietitian for help with recruitment and clinical advice; M. Lingaya (NIHR Nottingham Biomedical Research Centre) for preparation of stool samples and DNA extraction; D. Baker and R. Evans (Quadram Institute Biosciences) for DNA normalisation and sequencing library preparation; A. Wragg (NIHR Nottingham Biomedical Research Centre) for his assistance with the patient and public involvement group; C. Lam and S. Beg for their expert gastroenterology advice.

## CRediT Authorship Contributions

## Conflicts of interest

The authors declare no conflict of interest..

## Funding

CMC was supported by a College of Radiographers Doctoral Fellowship grant. The MRI scanning and other research costs were met by internal, academic research funds of the Nottingham Digestive Diseases Centre, University of Nottingham. The authors gratefully acknowledge the support of the Biotechnology and Biological Sciences Research Council (BBSRC); this research was funded by the BBSRC Institute Strategic Programme Food Microbiome and Health BB/X011054/1 and its constituent project BBS/E/F/000PR13631. This research was supported by the National Institute for Health and Care Research (NIHR) Nottingham Biomedical Research Centre. The views expressed are those of the authors and not necessarily those of the NHS, the NIHR, or the Department of Health & Social Care.

## Ethics approval

The patients underwent clinical tests and assessments as part of their routine care. This study involved human participants and ethics approval was obtained from the United Kingdom National Research Ethics Service (approval number 14MP002). The study was conducted in accordance with the Declaration of Helsinki (6th revision, 2008). All participants gave informed written consent to participate in the study before taking part.

## Data Availability

Raw read data from the metagenomic sequencing runs can be accessed through the NCBI SRA project number PRJNA1023737 and can be accessed at http://www.ncbi.nlm.nih.gov/bioproject/1023737. The associated metadata can be accessed through the University of Nottingham Research Repository DOI: 10.17639/nott.7352.

The full MATAFILER pipeline is available at https://github.com/hildebra/MATAF3. The RTK package is available at www.github.com/hildebra/Rarefaction/.

## Notes

http://www.ncbi.nlm.nih.gov/bioproject/1023737

### Competing Interest Statement

The authors have declared no competing interest.

## References

1. Rubio Tapia A. ACG Clinical Guidelines: Diagnosis and Management of Celiac Disease. Am J Gastroenterol. 2013;108:656–76.

2. Ciacci C, Ciclitira P, Hadjivassiliou M, et al. The gluten-free diet and its current application in coeliac disease and dermatitis herpetiformis. United European Gastroenterol J. 2015;3:121–35.

3. Midhagen G, Hallert C. High rate of gastrointestinal symptoms in celiac patients living on a gluten-free diet: controlled study. Am J Gastroenterol. 2003;98:2023–6.

4. Lam C, Sanders DS, Lanyon P, et al. Increased fasting small-bowel water content in untreated coeliac disease and scleroderma as assessed by magnetic resonance imaging. United European Gastroenterol J. 2019;7:1353–60.

5. Spiller RC, Lee YC, Edge C, et al. Delayed mouth-caecum transit of a lactulose labelled liquid test meal in patients with steatorrhoea caused by partially treated coeliac disease. Gut. 1987;28:1275-82.

6. Sadik R, Abrahamsson H, Kilander A, et al. Gut Transit in Celiac Disease: Delay of Small Bowel Transit and Acceleration after Dietary Treatment. Am J Gastroenterol. 2004;99:2429-36.

7. Pinto-Sanchez MI, Bercik P, Verdu EF. Motility alterations in celiac disease and non-celiac gluten sensitivity. Dig Dis. 2015;33:200-7.

8. Verdu EF, Galipeau HJ, Jabri B. Novel players in coeliac disease pathogenesis: role of the gut microbiota. Nat Rev Gastro Hepat. 2015;12:497-506.

9. Olivares M, Neef A, Castillejo G, et al. The HLA-DQ2 genotype selects for early intestinal microbiota composition in infants at high risk of developing coeliac disease. Gut. 2015;64:406-17.

10. de Meij TGJ, Budding AE, Grasman ME, et al. Composition and diversity of the duodenal mucosa-associated microbiome in children with untreated coeliac disease. Scand J Gastroenterol. 2013;48:530-6.

11. Ohlund K, Olsson C, Hernell O, et al. Dietary shortcomings in children on a gluten-free diet. J Hum Nutr Dietet 2010;23:294-300.

12. De Palma G, Nadal I, Collado MC, et al. Effects of a gluten-free diet on gut microbiota and immune function in healthy adult human subjects. Br J Nutr. 2009;102:1154-60.

13. Tjellstrom B, Stenhammar L, Hogberg L, et al. Gut microflora associated characteristics in children with celiac disease. Am J Gastroenterol. 2005;100:2784-8.

14. Biagi F, Andrealli A, Bianchi PI, et al. A gluten-free diet score to evaluate dietary compliance in patients with coeliac disease. Br J Nutr. 2009;102:882-7.

15. Chaddock G, Lam C, Hoad CL, et al. Novel MRI tests of orocecal transit time and whole gut transit time: studies in normal subjects. Neurogastroenterol Motil. 2014;26:205-14.

16. Segata N, Izard J, Waldron L, et al. Metagenomic biomarker discovery and explanation. Genome Biol. 2011;12.

17. Mallick H, Rahnavard A, McIver LJ, et al. Multivariable association discovery in population-scale meta-omics studies. PLoS Comput Biol. 2021;17.

18. Lewis SJ, Heaton KW. Stool form scale as a useful guide to intestinal transit time. Scand J Gastroenterol. 1997;32:920-4.

19. Prochazkova N, Falony G, Dragsted LO, et al. Advancing human gut microbiota research by considering gut transit time. Gut. 2023;72:180-91.

20. Asnicar F, Leeming ER, Dimidi E, et al. Blue poo: impact of gut transit time on the gut microbiome using a novel marker. Gut. 2021;70:1665-74.

21. Frioux C, Ansorge R, Özkurt E, et al. Enterosignatures define common bacterial guilds in the human gut microbiome. Cell Host Microbe. 2023;31:1111-25.

22. Song A-X, Li L-Q, Yin J-Y, et al. Mechanistic insights into the structure-dependant and strain-specific utilization of wheat arabinoxylan by Bifidobacterium longum. Carbohyd Polym. 2020;249:116886.

23. Nguyen NK, Deehan EC, Zhang Z, et al. Gut microbiota modulation with long-chain corn bran arabinoxylan in adults with overweight and obesity is linked to an individualized temporal increase in fecal propionate. Microbiome. 2020;8:1-21.

24. Wu X, Qian L, Liu K, et al. Gastrointestinal microbiome and gluten in celiac disease. Ann Med. 2021;53:1797-805.

25. Francavilla A, Ferrero G, Pardini B, et al. Gluten-free diet affects fecal small non-coding RNA profiles and microbiome composition in celiac disease supporting a host-gut microbiota crosstalk. Gut Microbes. 2023;15:2172955.

26. Caio G, Lungaro L, Segata N, et al. Effect of gluten-free diet on gut microbiota composition in patients with celiac disease and non-celiac gluten/wheat sensitivity. Nutrients. 2020;12:1832.

27. Hazard B, Trafford K, Lovegrove A, et al. Strategies to improve wheat for human health. Nat Food. 2020;1:475-80.

28. Leonard MM, Valitutti F, Karathia H, et al. Microbiome signatures of progression toward celiac disease onset in at-risk children in a longitudinal prospective cohort study. Proc Natl Acad Sci. 2021;118:e2020322118.

29. Smecuol E, Hwang HJ, Sugai E, et al. Exploratory, Randomized, Double-blind, placebo-controlled study on the effects of *Bifidobacterium infantis* natren life sart strain super strain in active celiac disease. J Clin Gastroenterol. 2013;47:139-47.

