## Supplementary Figure 1 for "One year of gluten free diet impacts gut function and microbiome in celiac disease"

Supplementary materials

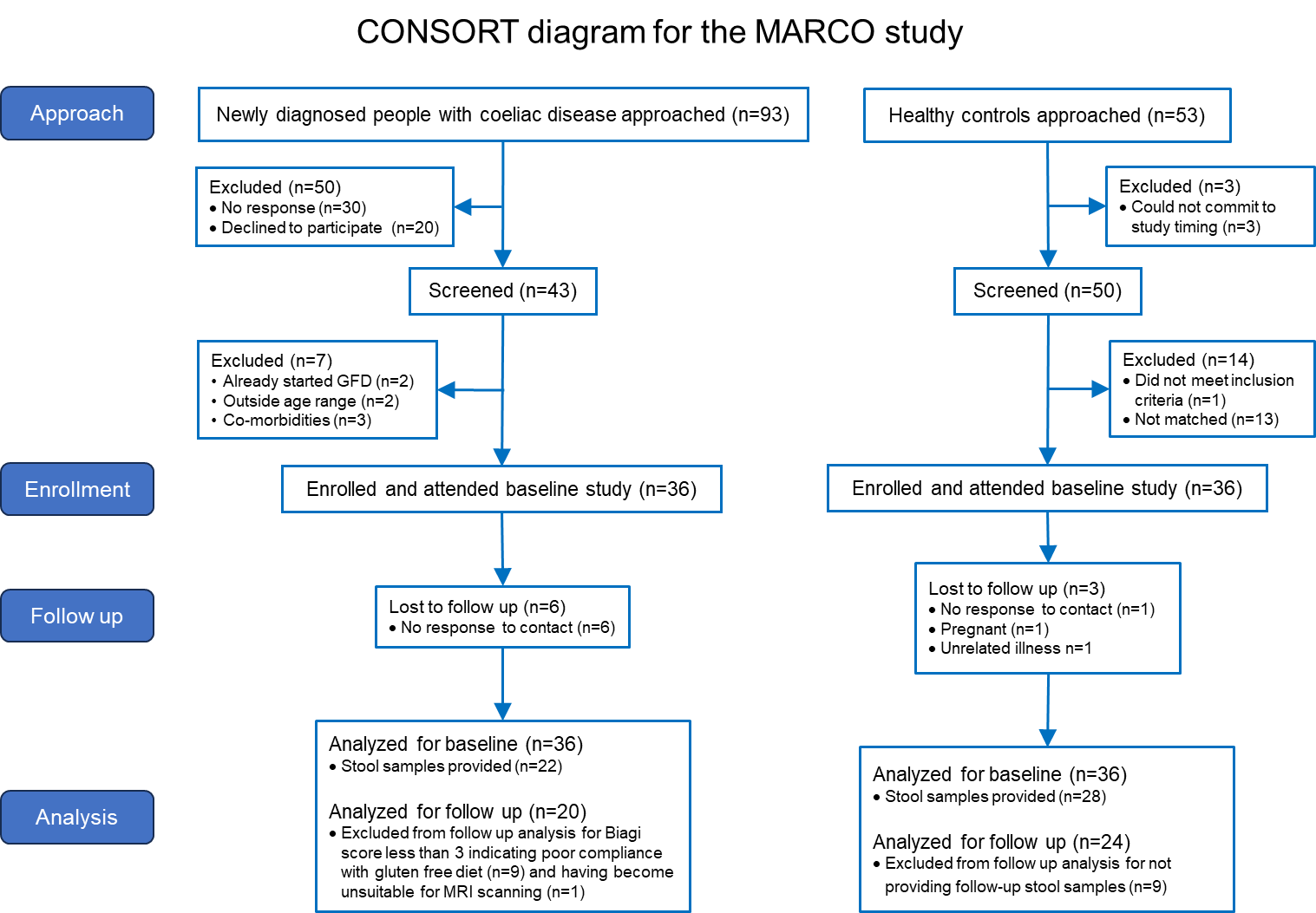

**Supplementary Figure 1.** CONSORT diagram for the study.

ONLINE SUPPLMENTAL MATERIALS AND METHODS

*Magnetic resonance imaging*

All participants were scanned supine on a 1.5T GE HDxt MRI scanner. The total scan duration was 15 minutes. After localising scans, three different imaging sequences were acquired using short breath-holds. A coronal T2 single shot fast spin echo sequence was used to measure the volumes of freely mobile SBWC. ^1^ This gave high‐intensity signal from areas with freely mobile fluid and little signal from body tissues. A coronal dual echo fast field echo sequence was then used to visualise the abdominal anatomy and measure colonic volumes. ^2^ Lastly, a coronal LAVA 3D fat saturated sequence was used to identify the transit markers. ^3^ To quantify whole gut transit time (WGTT), each transit capsule marker was allocated a score based on its position in the colon and WGTT was assessed using a weighted average position score (WAPS, in arbitrary units) of the five capsule markers.  ^3^ The WAPS can be extrapolated with some assumption to WGTT time in hours. ^3^ One of the patients’ MRI scans out of 20 and three of the controls’ MRI scans had poor quality due to respiratory motion and were excluded from the small bowel water content dataset.

*Short chain fatty acids*

The methods for faecal short chain fatty acid (SCFA) analysis were described previously. ^4^ Briefly, SCFAs were quantified by gas chromatography–mass spectrometry. Separation and detection of SCFAs of interest was achieved with splitless injection of the ethyl acetate extract using a Trace GC Ultra (Thermo Scientific, Manchester, UK) coupled with a DSQII mass spectrometer (Thermo Scientific). Compound identification was achieved by matching with database mass spectra (NIST/EPA/NIH Mass Spectral Library, Version 2.0d, NIST, Gaithersburg, MD, USA). Concentrations of analyte were calculated using ‘Xcalibur’ software (Thermo Scientific, UK).

*DNA extraction and metagenomic sequencing*

DNA was extracted using an adaptation of the Qiagen QIAmp DNA mini kit. Firstly, 0.125 g of defrosted stool sample was weighed into a 2 mL screw-cap microcentrifuge tube and was suspended in 0.5 mL of lysis buffer (500 mM NaCl,50 mM Tris-HCl (pH 8), 50 mM EDTA, 4 % SDS). The tubes were pre-loaded with 0.25 g of 0.1 mm zirconia beads and three 3 mm glass beads. The tubes were homogenised in a FastPrep for 1 min at 5.5 m/s then placed on ice for 30 s. This cycle was repeated two further times. The homogenised samples were heated to 95°C for 15 min, mixing by hand every 5 min. They were then cooled and centrifuged at 4°C for 5 min (13,000xg). The supernatant was transferred to a fresh 2 mL microcentrifuge tube. The pellet was resuspended in 150 µL of lysis buffer and the heating and centrifugation steps repeated, and the supernatants pooled. To each lysate tube, 130 µL of 10 M ammonium acetate was added, well mixed and incubated on ice for 5 min, before centrifuging (4°C, 13,000xg) for 10 min. The supernatant was then transferred to a fresh 2 mL microcentrifuge tube and an equal volume of isopropanol was added. This was mixed well and incubated on ice for 30 min. The tube was the centrifuged (4°C, 13,000 g) for 15 min, the supernatant discarded and the pellet washed with 0.5 mL of ethanol and then allowed to dry. The resulting pellet was then re-dissolved in 200 µL TE buffer, and 2 µL of DNase-free RNase (10 mg/mL) was added. The tube was then incubated at 37°C for 15 min, before adding 15 µL of proteinase K, 200µL of buffer AL and heating at 70°C for a further 10 min. After cooling to room temperature 200 µL of ethanol was added to the tube, and the contents were transferred to a QIAmp column. The QIAmp column was centrifuged at 6000xg for 1 minute, the flow through discarded and placed in a fresh collection tube. 500 µL of AW1 buffer was added to the column and then centrifuged at 6000 g for 1 minute, the flow through discarded and placed in a fresh collection tube. The column was then transferred to a fresh collection tube and 500 µL of AW2 buffer added. The column was then centrifuged at 20,000 g for 3 min, the flow-through discarded, and the column was then centrifuged for a further 1 min in a fresh collection tube to dry. The column was then placed in a fresh 1.5 mL microcentrifuge tube and 200 µL AE buffer added. This was then centrifuged at 6,000 g for 1 min to elute genomic DNA.

Genomic DNA was normalised to 5 ng/µL with elution buffer (10 mM Tris HCl). A miniaturised reaction was set up using the Nextera DNA Flex Library Prep Kit (Illumina, Cambridge, UK). 0.5 µL Tagmentation Buffer 1 (TB1) was mixed with 0.5 µL Bead-Linked Transposomes (BLT) and 4.0 µL PCR-grade water in a master mix and 5 µL was added to each well of a chilled 96-well plate. About 2 µL of normalised DNA (10 ng total) was pipette-mixed with each well of Tagmentation master mix and the plate heated to 55°C for 15 min in a PCR block. A PCR master mix was made up using 4 µL kapa2G buffer, 0.4 µL dNTP’s, 0.08 µL Polymerase and 4.52 µL PCR-grade water, from the Kap2G Robust PCR kit (Sigma-Aldrich, Gillingham, UK) and 9 µL added to each well in a 96-well plate. About 2 µL each of P7 and P5 of Nextera XT Index Kit v2 index primers (catalogue No. FC-131-2001 to 2004; Illumina, Cambridge, UK) were also added to each well. Finally, the 7 µL of Tagmentation mix was added and mixed. The PCR was run at 72 °C for 3 min, 95 °C for 1 min, 14 cycles of 95 °C for 10 s, 55 °C for 20 s and 72 °C for 3 min. Following the PCR reaction, the libraries from each sample were quantified using the methods described earlier and the high sensitivity Quant-iT dsDNA Assay Kit. Libraries were pooled following quantification in equal quantities. The final pool was double-SPRI size selected between 0.5 and 0.7X bead volumes using KAPA Pure Beads (Roche, Wilmington, US). The final pool was quantified on a Qubit 3.0 instrument and run on a D5000 ScreenTape (Agilent, Waldbronn, DE) using the Agilent Tapestation 4200 to calculate the final library pool molarity. qPCR was done on an Applied Biosystems StepOne Plus machine. Samples quantified were diluted 1 in 10,000. A PCR master mix was prepared using 10 µL KAPA SYBR FAST qPCR Master Mix (2X) (Sigma-Aldrich, Gillingham, UK), 0.4 µL ROX High, 0.4 µL 10 μM forward primer, 0.4 µL 10 μM reverse primer, 4 µL template DNA, 4.8 µL PCR-grade water. The PCR programme was: 95 °C for 3 min, 40 cycles of 95 °C for 10 s, 60 °C for 30 s. Standards were made from a 10 nM stock of Phix, diluted in PCR-grade water. The standard range was 20, 2, 0.2, 0.02, 0.002, 0.0002 pmol. The pooled library was then sent to Novogene (Cambridge, UK) for sequencing using an Illumina NovaSeq instrument, with sample names and index combinations used. Demultiplexed FASTQ’s were returned on a hard drive. A sequencing depth of ~10GB per sample was achieved.

*Metagenomic processing*

All metagenomic processing from raw reads to MGS abundance was conducted using the MATAFILER pipeline ^5^^,^ ^6^. Briefly, raw shotgun metagenomes were quality filtered using sdm v1.63 with default parameters ^7^, assembled using megahit v 1.2.9 with parameters “--k-list 25,43,67,87,101,127” ^8^, and reads mapped onto assemblies using bowtie2 v2.3.4.1 with parameters “--end-to-end “ ^9^, genes predicted with prodigal v2.6.1 with parameters “-p meta” ^10^ and a gene catalogue clustered at 95% nt identity using mmseqs2. ^11^ MAGs (metagenomic assembled genomes) were binned using SemiBin2 ^12^ and combined in MATAFILER to MGS (metagenomic species), relying on canopy clustering ^13^. Matrix operations were carried out using rtk ^14^.

To functionally annotate genes in the gene catalogue, dbCAN3 ^15^ was used to annotate genes to the CAZyme database and DIAMOND ^16^ to annotate KEGG orthologoues. Based on the KEGG Ortholog abundance matrices we further calculated KEGG and GMM ^17^ module abundances using a custom C++ implementation available on [www.github.com/hildebra/Rarefaction/](http://www.github.com/hildebra/Rarefaction/), similar to the protocol described in Forslund *et al*. ^18^.

The data were TSS normalised and log transformed prior to analysis. Corrected p-values were estimated using the Benjamini-Hochberg, BH, correction method. An FDR corrected p-value (q-value) of 0.1 was selected as a cut-off for significance. Due to the longitudinal nature of the dataset the stool donor was included in the model as a random effect. Abundance of the five Enterosignatures ^5^ were calculated using the web server <https://enterosignatures.quadram.ac.uk/>. PERMANOVA analysis was carried out between HV and CD microbiome at baseline and follow-up based on Bray-Curtis dissimilarity using the adonis2 function in the VEGAN library R package (v2.6.4). ^19^

When exploring associations with MRI variables, multivariate linear regression analyses were adjusted for multiple comparisons using Benjamini–Hochberg correction, with a significance threshold set to p=0.05, q = 0.1 using the MaAsLin2 software package.

*Patient and public involvement*

With the support an experienced Patient and Public Involvement & Engagement facilitator at the NIHR Nottingham Biomedical Research Centre and with the advice of charity Coeliac UK, a coeliac patient focus group was set up to inform the initial trial protocol. All aspects of the trial were discussed with members of this group. Their opinion on use of language and on the patient journey during the trial informed all patient-facing documentation and many practical aspects of the study e.g. the stool collection.

Supplementary References

1	Hoad CL, Marciani L, Foley S, Totman JJ, Wright J, Bush D, et al. Non-invasive quantification of small bowel water content by MRI: a validation study. Phys Med Biol 2007;52:6909-22.

2	Pritchard SE, Marciani L, Garsed KC, Hoad CL, Thongborisute W, Roberts E, et al. Fasting and postprandial volumes of the undisturbed colon: normal values and changes in diarrhea-predominant irritable bowel syndrome measured using serial MRI. Neurogastroenterol Motil 2014;26:124-30.

3	Chaddock G, Lam C, Hoad CL, Costigan C, Cox EF, Placidi E, et al. Novel MRI tests of orocecal transit time and whole gut transit time: studies in normal subjects. Neurogastroenterol Motil 2014;26:205-14.

4	Sloan TJ, Jalanka J, Major GAD, Krishnasamy S, Pritchard S, Abdelrazig S, et al. A low FODMAP diet is associated with changes in the microbiota and reduction in breath hydrogen but not colonic volume in healthy subjects. PLoS One 2018;13.

5	Frioux C, Ansorge R, Özkurt E, Nedjad CG, Fritscher J, Quince C, et al. Enterosignatures define common bacterial guilds in the human gut microbiome. Cell Host Microbe 2023.

6	Hildebrand F, Gossmann TI, Frioux C, Özkurt E, Myers PN, Ferretti P, et al. Dispersal strategies shape persistence and evolution of human gut bacteria. Cell Host Microbe 2021;29:1167-76. e9.

7	Özkurt E, Fritscher J, Soranzo N, Ng DY, Davey RP, Bahram M, et al. LotuS2: an ultrafast and highly accurate tool for amplicon sequencing analysis. Microbiome 2022;10:1-14.

8	Li D, Liu C-M, Luo R, Sadakane K, Lam T-W. MEGAHIT: an ultra-fast single-node solution for large and complex metagenomics assembly via succinct de Bruijn graph. Bioinformatics 2015;31:1674-6.

9	Langmead B, Salzberg SL. Fast gapped-read alignment with Bowtie 2. Nat Met 2012;9:357-9.

10	Hyatt D, Chen G-L, LoCascio PF, Land ML, Larimer FW, Hauser LJ. Prodigal: prokaryotic gene recognition and translation initiation site identification. BMC Bioinformatics 2010;11:1-11.

11	Steinegger M, Söding J. MMseqs2 enables sensitive protein sequence searching for the analysis of massive data sets. Nat Biotechnol 2017;35:1026-8.

12	Pan S, Zhao X-M, Coelho LP. SemiBin2: self-supervised contrastive learning leads to better MAGs for short-and long-read sequencing. bioRxiv 2023:2023.01. 09.523201.

13	Nielsen HB, Almeida M, Juncker AS, Rasmussen S, Li J, Sunagawa S, et al. Identification and assembly of genomes and genetic elements in complex metagenomic samples without using reference genomes. Nat Biotechnol 2014;32:822-8.

14	Saary P, Forslund K, Bork P, Hildebrand F. RTK: efficient rarefaction analysis of large datasets. Bioinformatics 2017;33:2594-5.

15	Zheng J, Ge Q, Yan Y, Zhang X, Huang L, Yin Y. dbCAN3: automated carbohydrate-active enzyme and substrate annotation. Nucleic Acids Research 2023:gkad328.

16	Buchfink B, Xie C, Huson DH. Fast and sensitive protein alignment using DIAMOND. Nat Met 2015;12:59-60.

17	Darzi Y, Falony G, Vieira-Silva S, Raes J. Towards biome-specific analysis of meta-omics data. ISME J 2016;10:1025-8.

18	Forslund K, Hildebrand F, Nielsen T, Falony G, Le Chatelier E, Sunagawa S, et al. Disentangling type 2 diabetes and metformin treatment signatures in the human gut microbiota. Nat 2015;528:262-6.

19	Dixon P. VEGAN, a package of R functions for community ecology. J Veg Sci 2003;14:927-30.

Symptoms in patients with CD

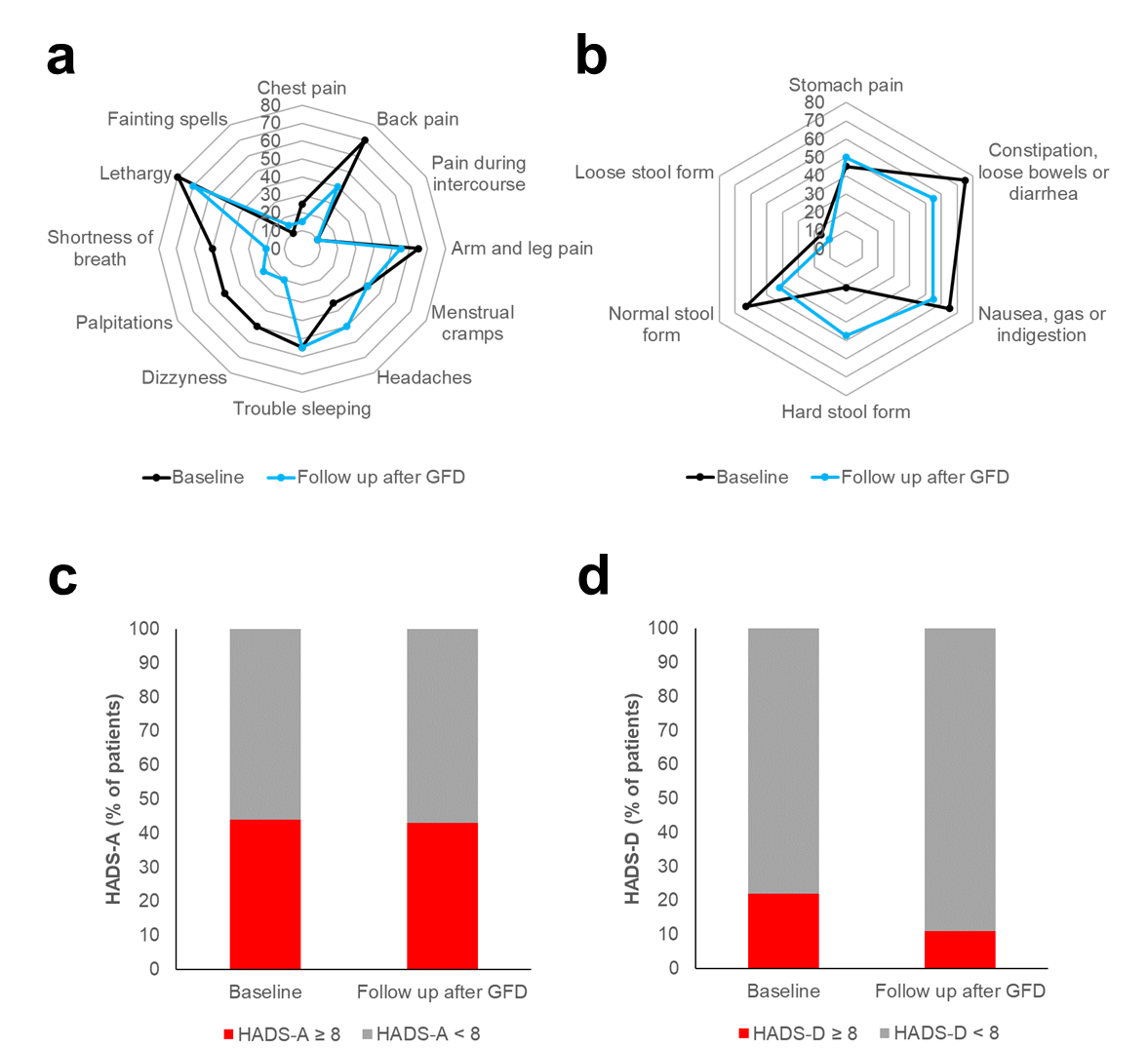

**Supplementary Figure 2.** Symptoms in patients with coeliac disease and changes following 12 months of gluten-free diet. n=20. People newly diagnosed with coeliac disease were studied before starting gluten-free diet treatment and 12 months after. At these two time points they filled in the Patient health Questionnaire (PHQ-15), the Hospital Anxiety and Depression Scale (HADS) and the Bristol Stool Form Scale (BSFS). (a) Radar diagram of the 12 non-gastrointestinal somatization symptoms of the PHQ questionnaire, before and after following the GFD. Each data point shows the percentage of patients ‘experiencing the symptom, corresponding to a score of 1 or 2 on the PHQ questionnaire for that symptom. (b) Radar diagram of the 3 gastrointestinal somatization symptoms of the PHQ questionnaire, before and after following the GFD. The diagram also shows form of the patients’ stools from the BSFS scale, whereby a score of 1-2 indicates hard stool form, a score of 3-4-5 indicates normal stool form and a score of 6-7 indicates loose stool form. Each data point shows the percentage of patients experiencing either the symptom, corresponding to a score of 1 or 2 on the PHQ questionnaire for that symptom or their stool form. (c) Bar chart of the anxiety subscale of the HADS questionnaire. A score of 8 or greater is often used to indicate clinical levels of anxiety (Bjelland et al. Journal of Psychosomatic Research 2002, 52, 69– 77). The chart shows the percentage of patients with clinical levels of anxiety before and after 1 year of GFD. (d) Bar chart of the depression subscale of the HADS questionnaire. A score of 8 or greater is often used to indicate clinical levels of depression (Bjelland et al. Journal of Psychosomatic Research 2002, 52, 69– 77). The chart shows the percentage of patients with clinical levels of depression before and after 1 year of GFD.

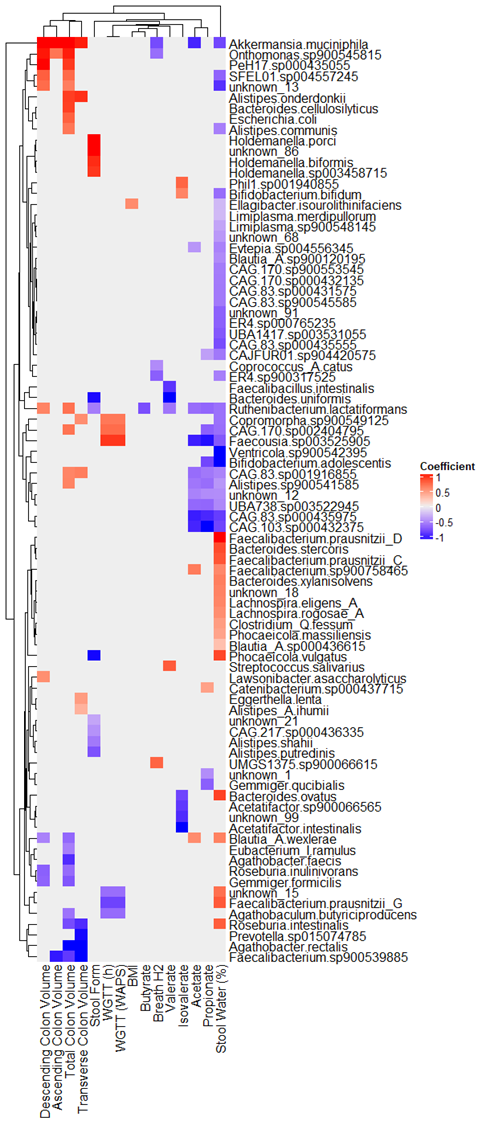

**Supplementary Figure 3.** Associations between MRI parameters, stool metabolites and water content, breath hydrogen, BMI, symptom scores (HADS and PHQ) and species identified by metagenomic sequencing. Associations were identified using the generalised linear modelling method in MaAsLin2 with TSS normalisation and log transformation of data. Only significant associations (q-value = <0.1) are shown, with the colour indicating a positive association (red) or a negative association (blue)

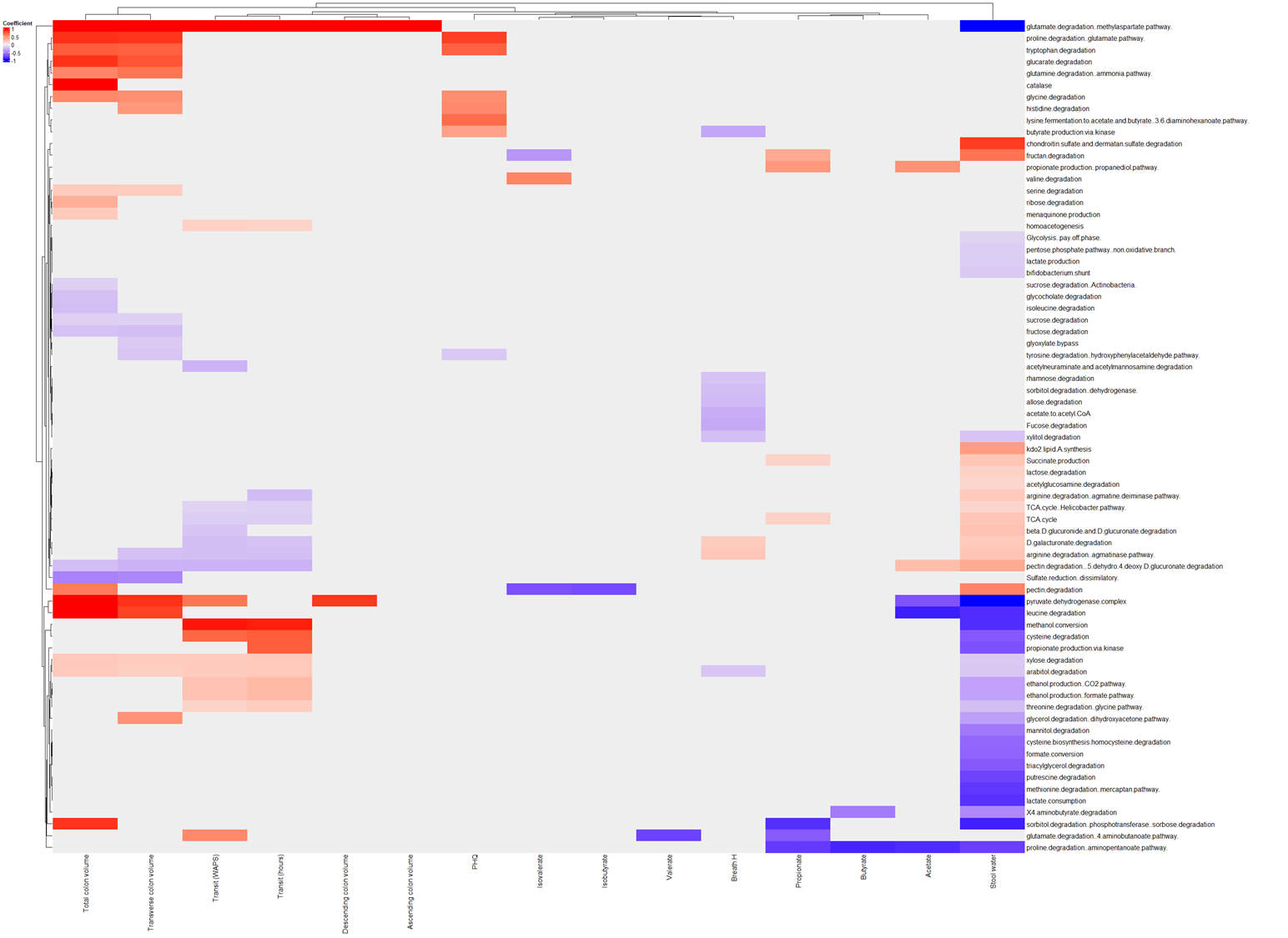

**Supplementary Figure 4.** Associations between MRI parameters, stool metabolites and water content, breath hydrogen, BMI, symptom scores (HADS and PHQ) and microbial metabolic pathways. Associations were identified using the generalised linear modelling method in MaAsLin2 with TSS normalisation and log transformation of data. Only significant associations (q-value = <0.1) are shown, with the colour indicating a positive association (red) or a negative association (blue).

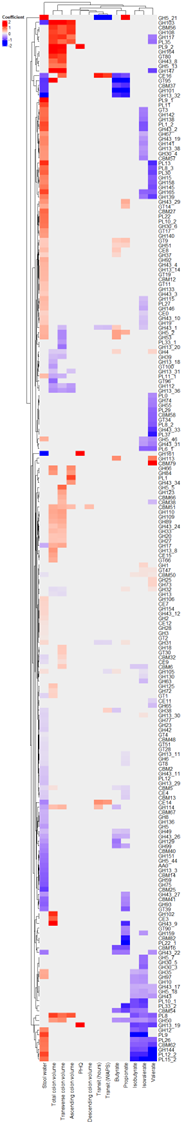

**Supplementary Figure 5.** Associations between MRI parameters, stool metabolites and water content, breath hydrogen, BMI, symptom scores (HADS and PHQ) and CAZyme family abundances. Associations were identified using the generalised linear modelling method in MaAsLin2 with TSS normalisation and log transformation of data. Only significant associations (q-value = <0.1) are shown, with the colour indicating a positive association (red) or a negative association (blue).

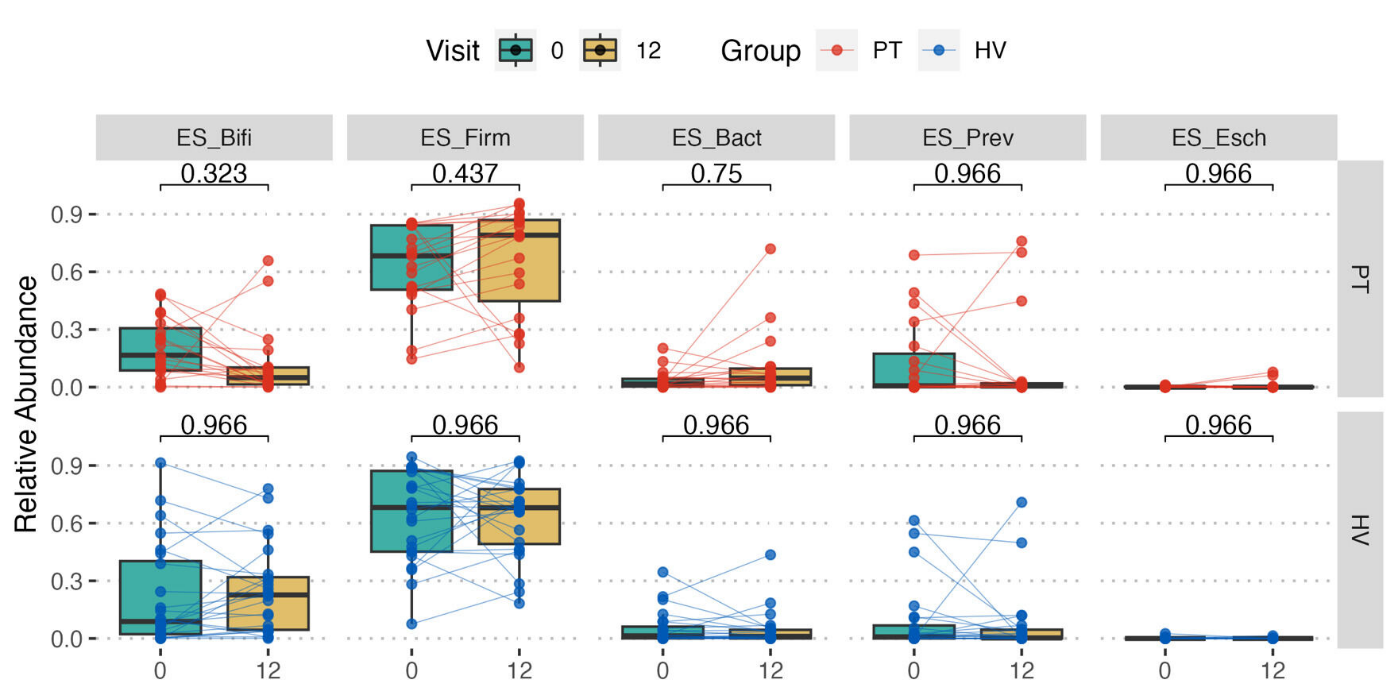

**Supplementary Figure 6.** Shifts in enterosignatures from baseline to follow-up in the HV and PT groups. The statistical significance of associations between enterosignature groups between baseline and follow-up for each group was calculated using a paired Wilcoxon test with Benjamini-Hochberg correction for multiple comparisons.

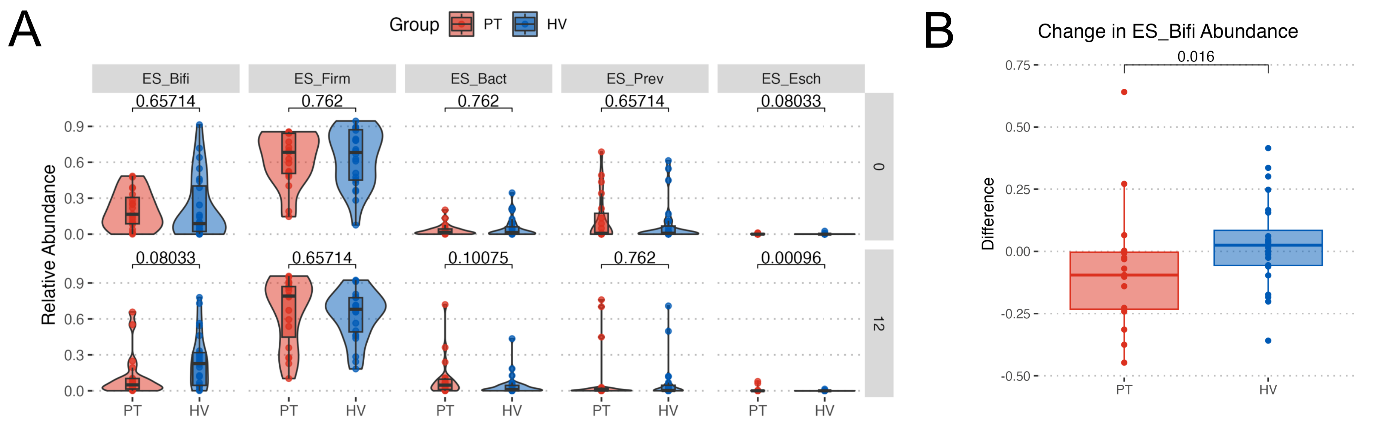

**Supplementary Figure 7.** Differences between enterosignatures between healthy volunteers (HV, n=24) and patients with coeliac disease (PT, n=20) at baseline and following 12 months of follow-up. (A) Shifts in relative abundance of enterosignatures. Statistical significance is indicated (q-values) on the basis of unpaired Wilcoxon tests with Benjamini-Hochberg correction for multiple comparisons. (B) Differential abundance of the ES_Bifi enterosignature from baseline to follow-up. Statistical significance is indicated on the basis of an unpaired Wilcoxon test.

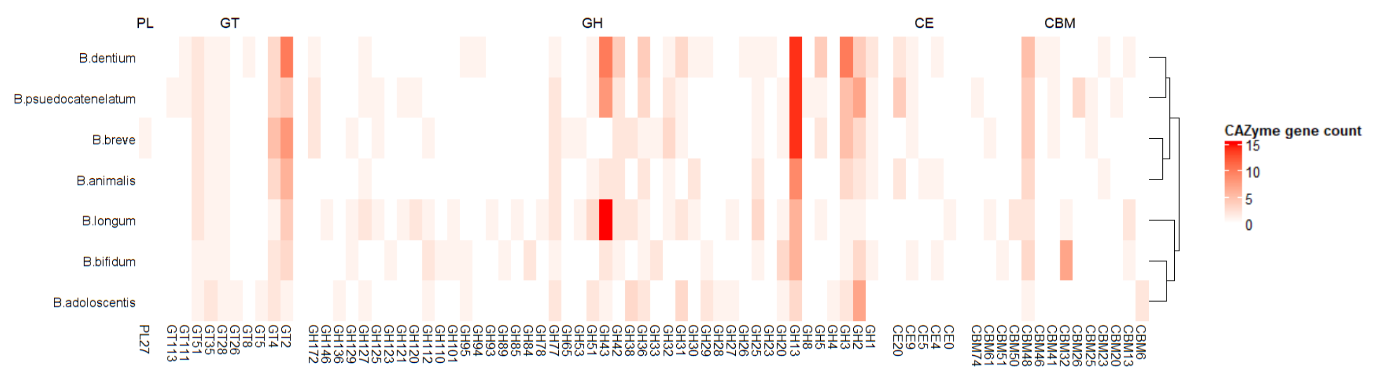

**Supplementary Figure 8.** CAZyme gene counts in representative metagenome assembled genomes of *Bifidobacterium* species identified in the study. PL = Polysaccharide Lyase, GT = Gycosyl Transferase, GH = Glycosyl Hydrolase, CE = Carbohydrate Esterase and CBM = Carbohydrate Binding Module. CAZyme counts were determined from the DBcan algorithm based on the CAZy database. Hits were only considered significant if identified by all three tools in the algorithm (DIAMOND, HMMER and dbCAN-sub).
